# De-heterogeneity of the eukaryotic viral reference database (EVRD) improves the accuracy and efficiency of viromic analysis

**DOI:** 10.1101/2022.03.03.482774

**Authors:** Junjie Chen, Xiaomin Yan, Yue Sun, Zilin Ren, Guangzhi Yan, Guoshuai Wang, Yuhang Liu, Zihan Zhao, Yang Liu, Changchun Tu, Biao He

**Affiliations:** Changchun Veterinary Research Institute, Chinese Academy of Agricultural Sciences, Changchun, Jilin Province, China; Jiangsu Co-innovation Center for Prevention and Control of Important Animal Infectious Diseases and Zoonosis, Yangzhou University, Yangzhou, Jiangsu Province, China

**Keywords:** Eukaryotic virome, emerging infectious disease, database contamination, host contamination, heterogenous sequences

## Abstract

Widespread in public databases, the notorious contamination in virus reference databases often leads to confusing even wrong conclusions in applications like viral disease diagnosis and viromic analysis, highlighting the need of a high-quality database. Here, we report the comprehensive scrutiny and the purification of the largest viral sequence collections of GenBank and UniProt by detection and characterization of heterogeneous sequences (HGSs). A total of 766 nucleotide- and 276 amino acid-HGSs were determined with length up to 6,605 bp, which were widely distributed in 39 families, with many involving highly public health-related viruses, such as hepatitis C virus, Crimea-Congo hemorrhagic fever virus and filovirus. Majority of these HGSs are sequences of a wide range of hosts including humans, with the rest resulting from vectors, misclassification and laboratory components. However, these HGSs cannot be simply considered as exotic contaminants, since part of which are resultants of natural occurrence or artificial engineering of the viruses. Nevertheless, they significantly disturb the genomic analysis, and hence were deleted from the database. A further augmentation was implemented with addition of the risk and vaccine sequences, which finally results in a high-quality eukaryotic virus reference database (EVRD). EVRD showed higher accuracy and less time-consuming without coverage compromise by reducing false positives than other integrated databases in viromic analysis. EVRD is freely accessible with favorable application in viral disease diagnosis, taxonomic clustering, viromic analysis and novel virus detection.

## Background

Emerging infectious diseases (EIDs), especially the viral ones, are a serious threat to public health, significantly challenging global security, social economy and human’s life (1). Rapid and accurate diagnosis of EIDs is a prerequisite for timely formulating and implementing prevention and control measures. High-throughput sequencing (HTS)-based metagenomics is a promising approach for rapid diagnosis and identification of EIDs because it does not require ‘*a priori*’ information and is capable of identifying a comprehensive spectrum of potential agents, especially those new ones, by a single test (2, 3). Metagenomic diagnosis highly depends on the similarity-based analyses of reads or contigs against reference database. Hence, the high quality of reference database that is of complete representativeness, functional robustness, and informational accuracy provides an important guarantee of diagnostic reliability.

There are numerous resources focusing on particular viruses. The Hepatitis B virus database (HBVdb) is a nucleotide (nt) and amino acid (aa) sequence collection for surveillance of genetic variability and analysis of drug resistance profiling of HBV (4). The HIV, HCV and HFV/Ebola databases incorporated in the Pathogen Research Databases contain data on viral genetic sequences, immunological epitopes and vaccine trials (https://www.lanl.gov/collaboration.pathogen-database/index.php). The Global Initiative on Sharing All Influence Data (GISAID) initially archived genetic sequences and related clinical and epidemiological data of all influenza viruses, and now has expanded to include the coronavirus causing COVID-19 (https://www.gisaid.org). Besides, several comprehensive databases covering a broad range of, even all, viruses have been established. The Virus Pathogen Databases and Analysis Resource (ViPR) provides cross-referenced data of multiple types on all high priority human pathogenic viruses (5). The Databases of Bat- and Rodent-associated Viruses (DBatVir and DRodVir) catalog all viral sequences discovered from the two most important viral natural hosts (6, 7). As the largest public biological sequence database, GenBank contains the viral and phage divisions that are widely used for genomic analysis (8). Similarly, the taxon Viruses of the UniProt knowledgebase (UniProtKB) provides a comprehensive set of viral protein sequences (9). The Reference Viral Database (RVDB) and its protein counterpart, RVDB-prot, were established to include all viral, virus-related, and virus-like entries (10, 11). The Integrated Microbial Genomes/Virus (IMG/VR) provides access to the largest collection of viral sequences obtained from (meta)genomes, among which more than 90% are bacteriophage (12).

These specialized databases focus on a taxonomic group or type of viruses, making them less representative. These comprehensive resources contain a high degree of redundancy. Of particular importance is that there are notable levels of heterogenous sequences (HGSs) in those databases (13). We define a sequence as heterogenous if it has a real identity inconsistent with its definition or is an exotic contaminant. Based on our experiences of viromic studies over the past decade, those HGSs are mainly related to laboratory components and nonviral organisms or artefacts. The laboratory component-derived sequences (LCDs), such as those of parvovirus-like hybrid virus (14), xenotropic murine leukemia virus-related virus (15) and human endogenous retrovirus H (16), are technically viral, but often carried by nucleic acid extraction spin columns, biologicals or experimental performers, and are very easy to contaminate samples, resulting in wrong conclusion in analyses (14-18). For example, parvovirus was erroneously diagnosed in dairy cattle with fever and diarrhea, but which was found to be a contaminant originating from Qiagen extraction columns (17). The nonviral sequences are actual artefacts derived from vectors or other organisms, but are misannotated as virus in reference databases, which are particularly problematic for viromic studies, in that if a genomic fragment of nonviral organism labeled as virus in a database, any samples from the organism might erroneously be determined to contain the virus. These HGSs are often inserted into large DNA viruses (LDVs) with most related to eukaryotic microorganisms or aquatic samples, e.g., mimivirus, pandoravirus and phycodnavirus. In animal viromic studies, a large number of sequences can be annotated to LDVs, even using a very stringent criterion. But most of those sequences were finally proven to originate from hosts, bacteria or other organisms. Some LDVs, such as herpesviruses, can integrate their genomic fragments into host genomes (19, 20), and viral genomes may also be misassembled to contain pieces of host sequences that are erroneously annotated as virus in database. In both cases, those Trojan horse-like sequences will greatly increase false positives in viromic analysis. These issues are very prone to draw a questionable even wrong conclusion and pose a great obstacle in applications like EID diagnosis, taxonomic classification and viromic studies, etc. (17, 18, 21-23), highlighting the need of a high-quality reference database.

To address these issues, here we established a stringent scrutiny pipeline to systematically analyze and identify HGSs concealed in the largest viral nt (GenBank) and aa (UniProt) reference collections, resulting in a nonredundant and well-refined eukaryotic viral reference database. To augment its function for diagnosis, we incorporated risk and vaccine information into the database, which helps identify possible exotic contamination and distinguish vaccine strains from field viruses. The database is expected to provide a more accurate reference for EID diagnosis, new virus identification, viromic analysis, and other virologic studies.

## Results and Discussion

### Overview of heterogenous sequences

The viral division (gbvrl) of GenBank is the largest resource of eukaryotic viral sequences, and widely used in virologic research, even construction of specialized sub-databases (5, 10), from which the Viral Genome Resources is derived to serve as a set of high-quality curated viral reference genomes and their validated genomic neighbors, but lacking the full-spectrum of viral diversity (24). As of March 04 2021, gbvrl and the Viral Genome Resources have archived 3,316,373 and 288,226 nt sequences, respectively. They overlapped 263,895 sequences, hence we added the remaining 24,331 sequences of the Viral Genome Resources into gbvrl, which brought to a preliminary data set (PDS) of 3,340,704 sequences. This data set was subjected to a stringent heterogeneity scrutiny pipeline, which is composed of five parts, i.e., preliminary filtration, host genome scrutiny, vector sequence scrutiny, annotation cross scrutiny, and cross check of viral metagenomes (Methods). Since we aimed to build a refined reference database for diagnosis of viral diseases and discovery of eukaryotic viruses, hence the first preliminary filtration step removed 91,549 sequences of viruses infecting bacteria, archaea, fungi or microorganisms, or shorter than 200 bp. After four rounds of scrutiny, we further removed and trimmed 146 and 373 sequences, respectively, with detection of 766 HGSs (some sequences have multiple HGSs).

These HGSs came from 39 viral families and unclassified viruses at the family level, with majority being *Herpesviridae* (59.9%), followed by *Flaviviridae* (14.0%) (Fig. 1). They were either full-length sequences (14.5%) or just chimeric fragments (85.5%) within viral genomes, and could be classified into four origins, i.e., host, vector, cross-host, and cross-family (Fig. 1), which likely originated from hosts and vectors, simultaneously appeared in viromic data of different hosts, and are misclassified at the family level, respectively. Their submission could be traced back to 1993 with 66.2% from 2015-2019 (Fig. 1). HTS-based viral metagenomics has dramatically expanded the space of our known viral sequences (25), but with an annoying side-effect, i.e., the chimeric viral assembly containing insertion of other viral sequences even sequences of other organisms (26). Though a lot of HGSs did not provide the information of sequencing technology in GenBank, we did find a substantial number of host HGSs (n>51) submitted since 2015 are probably due to the *de novo* assembly of Illumina reads. Majority (80.7%) of these HGSs were ≤ 600 bp, with a few within the families of *Papillomaviridae* (n=3), *Paramyxoviridae* (n=1), *Flaviviridae* (n=1) and *Herpesviridae* (n=3) exceeding 2,000 bp, even one HGS of *Herpesviridae* reaching 6,605 bp, all of which were host-origin with the exception of the *Paramyxoviridae* HGS that was related to vector (Fig. 1).

**Fig 1.**
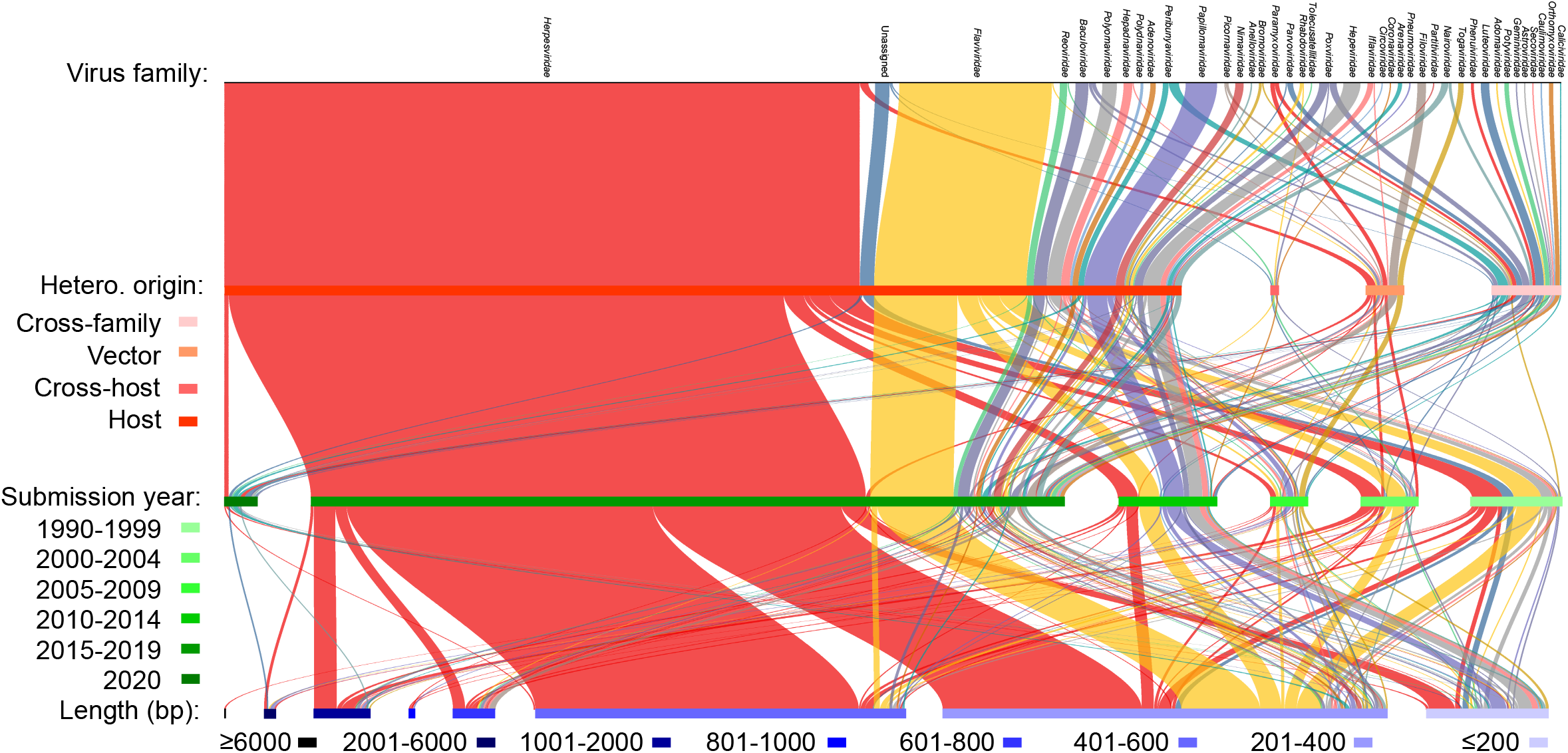
Summary of the 766 HGSs. Representing 39 viral families, they were classified into heterogeneity origins (Hetero. origin) of cross-family, vector, cross-host and host, with submission years to GenBank of 1993-2021 and length up to 6,605 bp.

Regarding aa sequences, we retrieved all sequences under the Taxonomy of Viruses in UniProtKB (version 2021_03). UniProtKB is mainly based on the translation of genome sequence submitted to the International Nucleotide Sequences Database Collaboration (INSDC) source databases, and also supplemented by genomes sequenced and/or annotated by other academic groups, making it as the most comprehensive set of protein sequences (9). Generally, UniProt aa sequences showed less heterogeneity compared to GenBank nt sequences, in that translation itself is a recognized validation means of viral genomes, and furthermore, heterogenous insertion often occurs as a flanking sequence in the untranslated region at the terminus of nt sequence. Finally, a total of 267 HGSs were detected with most being counterparts in nt scrutiny, hence which will not be discussed in details herein after.

### Various origins of HGSs and their causation: natural vs artificial

Among the four origins of HGSs, host sequences were predominant (86.9%), and were detected in 24 viral families (unclassified viruses were not counted) (Fig. 1). These host HGSs were related to humans and other animals covering non-human primates, bovines, canines, avians, rodents, bats and arthropod, etc., and even bacteria. HGSs within different families are prone to be dominated by certain heterogeneity types, e.g., almost all HGSs within *Herpesviridae* (96.3%) and *Flaviviridae* (99.1%) were associated with host genomes, while those *Togaviridae* and *Filoviridae* HGSs were all vector sequences (Fig. 1).

Heterogeneity is widespread in nonviral databases, in which human sequences were usually found to contaminate the genomic databases of bacteria, plants and fish, therefore those HGSs were all considered contaminants (27, 28). Merchant *et al*. found microbial sequences in cow genomes, but the final verification indicated that such contamination was due to that multiple sequences of *Neisseria gonorrhoeae* were actually derived from the cow or sheep genomes (29). Notably, a large-scale search has identified contamination of more than 2,000,000 exogenous sequences in the RefSeq, GenBank, and nr databases (13). However, we found that these viral HGSs cannot be simply considered contaminants, and can be classified as natural, intentionally artificial (ia) and unintentionally artificial (ua) ones based on their causation.

#### Natural heterogeneity

Some HGSs are naturally acquired by viruses in the process of proliferation, which are essential for certain viruses to gain new features. Bovine viral diarrhea virus (BVDV) is a worldwide distributing pathogen and can cause severe consequences to cattle and sheep (30). Almost all HGSs within the family *Flaviviridae* are inserts of bovine hybrid ribosomal S27a and ubiquitin sequences into the BVDV genomes (Fig. 2A). The in-frame insertion of the host sequence into NS3 gene is essential for the virus to gain cytopathogenicity in cell culture (31). Hepatitis E virus (HEV) is hardly cultured using cell systems, the integration of a short piece of human S17 ribosomal protein fragment into the hypervariable region of HEV genome (accession number: JQ679013) enables some variants to grow in HepG2/C3A cells (32).

**Fig 2.**
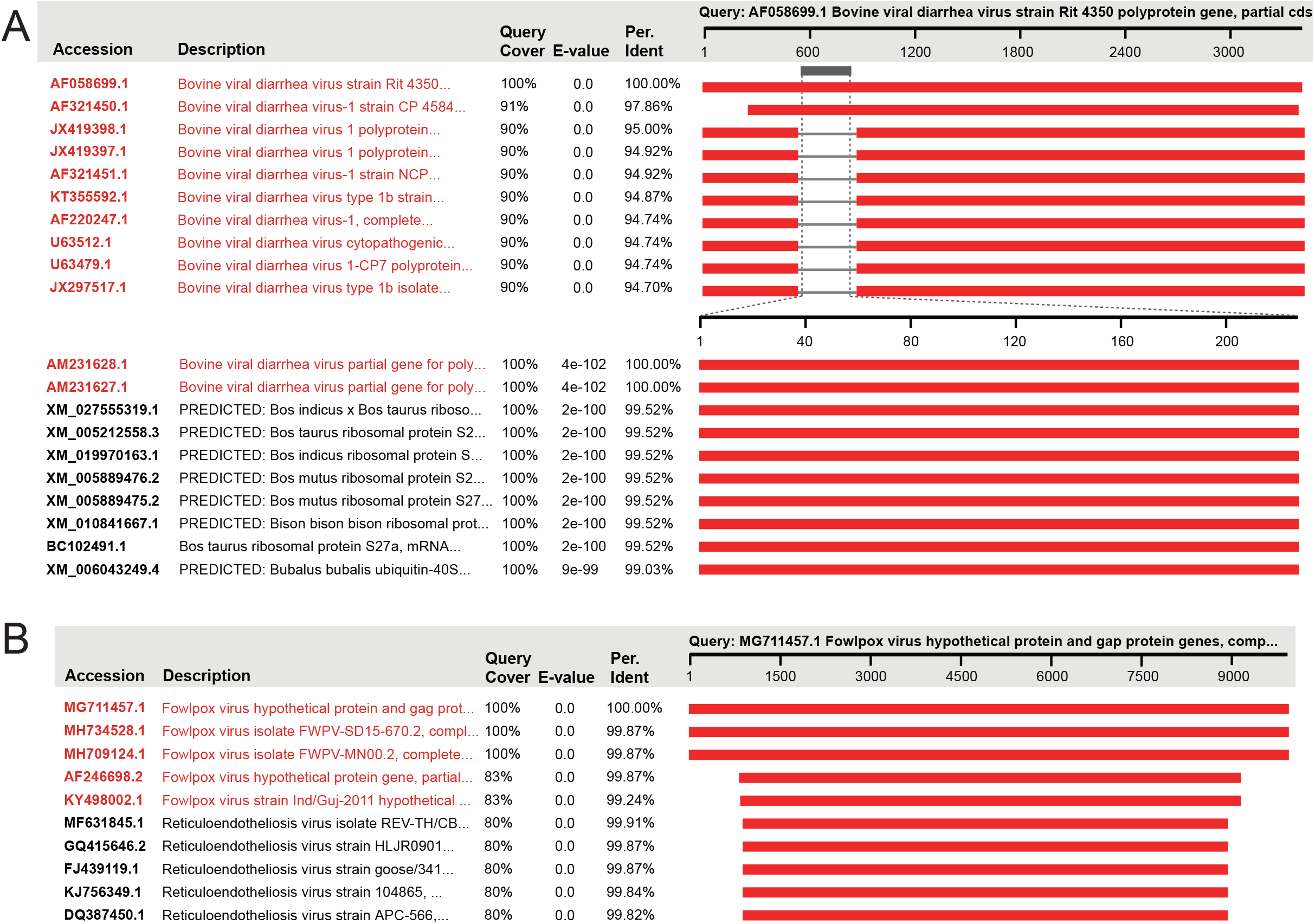
Identification of the naturally occurred HGSs of BVDV (A) and fowlpox virus (B) using blastn search. The blastn hits with close definition to the query are highlighted in red.

Besides host sequences, genomic fragments of other viral families can also integrate into some viral genomes, particularly during coinfection of multiple viruses. For some LDVs, viral DNA replicates within the cellular nucleus or cytoplasm, providing an opportunity for viral genome to be integrated by retrovirus. Thus avian retrovirus was shown to be integrated into the genome of Marek’s disease virus, an avian herpesvirus (33). We also detected reticuloendotheliosis viral sequences of various length, even near-full-length, integrated into genomes of some fowpox viruses (Fig. 2B), which likely enhanced the pathogenic trait of the virus (34, 35). Inter-family recombination can also occur in RNA viruses. A betacoronavirus detected in bats contained a unique gene integrated into the 3’-end of its genome that most likely originated from the p10 gene of a bat orthoreovirus, a gene that can induce the formation of cell syncytia (36).

#### Intentionally artificial heterogeneity

Some viral genomes are intentionally engineered to contain HGSs that might derive from nonviral artefacts or viruses of different families, by which these engineered viruses were used to study viral infection, deliver heterogenous proteins, even combat viral infectious diseases. We found that a large part of vector-(87.2%) and a few cross-family-(n=3), but no host HGSs are intentionally artificial. Among ia-vector HGSs, green fluorescent proteins are very common (41.5%) (Fig. 3A), and elements like neomycin phosphotransferase, mCherry and firefly luciferase can also be observed. The three ia-cross-family HGSs are all associated with avian paramyxovirus within the family *Paramyxoviridae*. These recombinants were generated using reverse genetics to serve as vaccine vector expressing the hemagglutinin of highly pathogenic avian influenza virus to induce protective immunity against influenza virus in chickens (Fig. 3B) (37).

**Fig 3.**
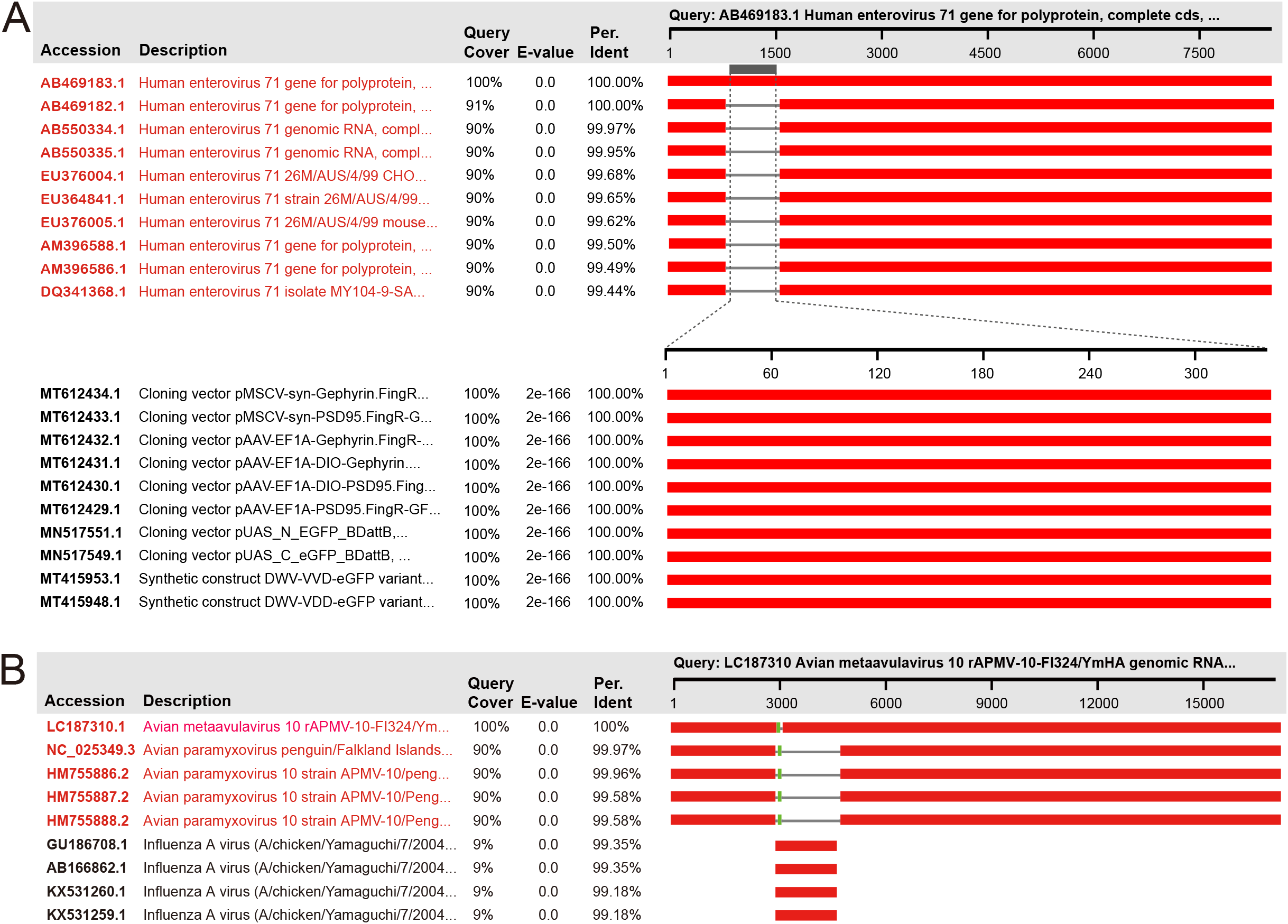
Identification of the ia-HGSs of human enterovirus 71 (A) and avain metaavulavirus (B) using blastn search. The blastn hits with close definition to the query are highlighted in red.

#### Unintentionally artificial heterogeneity

The ua-HGSs are technically true errors, but are unintentionally annotated as viral components. They are widely distributed in host-, vector-, cross-host- and cross-family-HGSs. The ua-host HGSs can be full-length sequences, e.g., a 399 bp-long human mRNA was erroneously defined hepatitis C virus (Fig. 4A). *de novo* assembly of HTS reads occasionally results in chimeric ua-host HGSs often at the termini of sequence, e.g., a 1,636 bp-long human sorting nexin 10 fragment was misassembled into the 3’ terminus of the segment M of a Crimean-Congo hemorrhagic fever orthonairovirus (CCHFV) (Fig. 4B). As to ua-vector HGSs, we found two short stealth virus sequences that are actually vector backbones. Through cross check of viral metagenomes from different hosts, we found 5 commonly existing HGSs, which shared >99% nt identities with the sequences in viromic data of different host species. Viruses harbored by different host species usually show significant genetic distances. If a virus is found in hosts of different highly-hierarchic taxon, it should be noted whether it results from cross-species transmission or just contamination. Further verification showed that the five references are all non-viral, but genomic fragments of bacteria. For example, a blue tongue virus reference (AY397620) frequently found in our viral metagenomic analyses is a *Mycoplasma bovis* chromosomic sequence.

**Fig 4.**
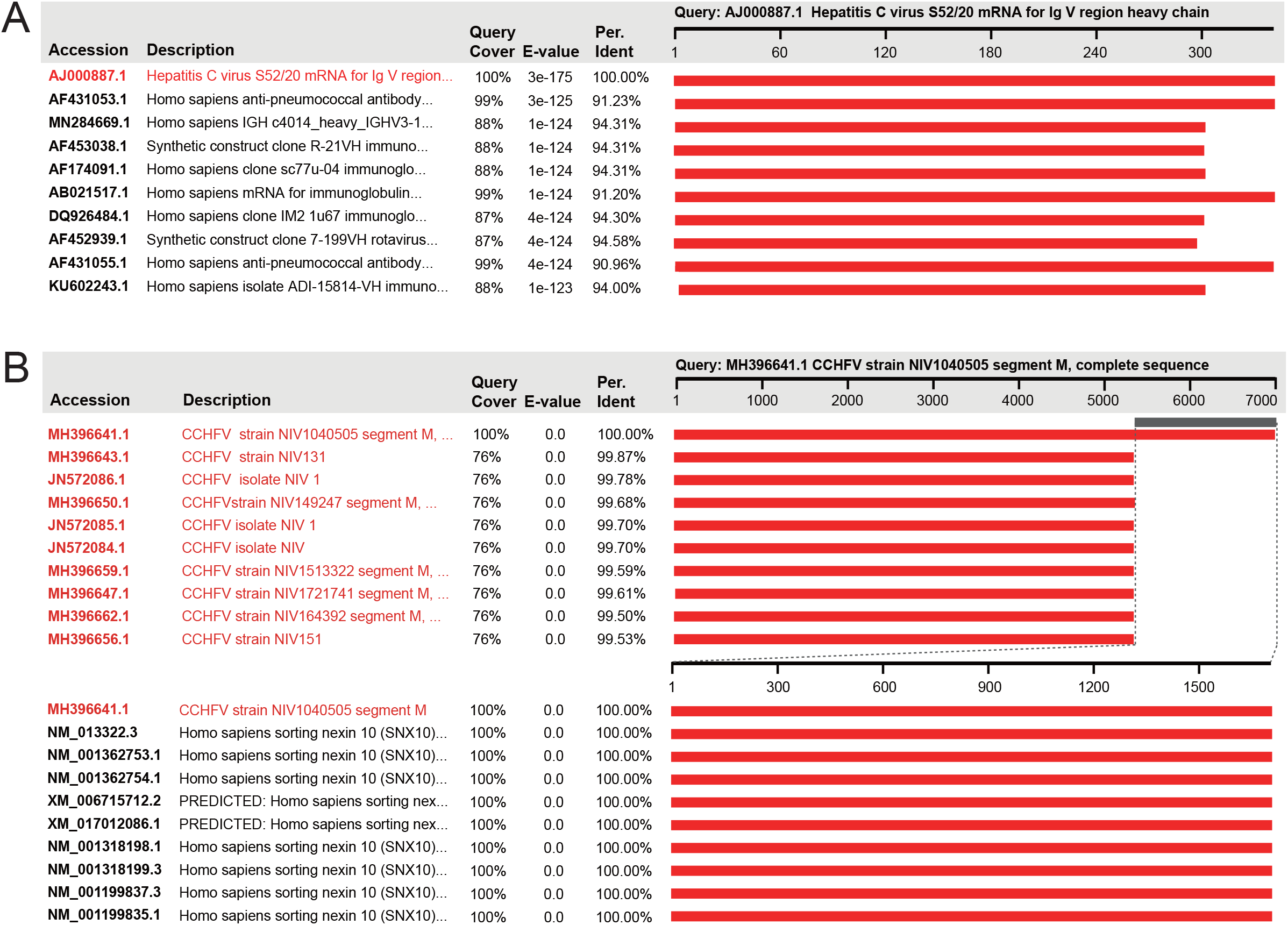
Identification of the ua-HGSs of hepatitis C virus (A) and CCHFV (B) using blastn search. The blastn hits with close definition to the query are highlighted in red.

Cross-family misclassification can occur between eukaryotic viral families, even between eukaryotic and prokaryotic viral families. Three sequences wrapping circovirus-featured *rep* and *cap* genes should be classified into the family *Circoviridae*, but are defined dependoparvoviruses within the family *Parvoviridae*. A 558 bp-long sapovirus sequence (AB212270) defined within the family *Caliciviridae* actually originated from bacterophage since it has almost all high-quality nt and aa blast hits against Salmonella phages. If a viral sequence is highly novel with very low identity to known references, it would be misclassified at the family level. A 4,047 bp-long sequence recovered from a bird metagenome was defined *Parvoviridae sp*., but which had very few blastn hits in nt database and several blastx hits against major capsid proteins of microviruses. Profile comparison showed that, though with very low identity and similarity, one of its encoding products perfectly matched to the capsid protein of microvirus, a viral hallmark gene, with probability of 100%. Accordingly, it should be classified as a bacteriophage than a parvovirus.

### Augmentation by adding warning sequences

Though these natural and ia-HGSs endow viruses with some necessary functions, and are not so-called contaminants. They do result in heterogeneity to viral genomes, along with ua-HGSs, which are substantially problematic in virus identification, viromic annotation and taxonomic assignment. To establish a neat reference database, we deleted the HGSs to minimize the heterogeneity of existing reference database. However, the resulted database is still redundant with high level of identical sequences. Thus, a de-redundance at 99% identity and 90% coverage was conducted, which downsized the nt and aa databases for ∼6 and ∼3 times, respectively.

Augmentation was implemented to the database with addition of tagged LCD (n=155), viral functional cassette (n=79) and vaccine (n=40) sequences to the nt reference database. The LCD sequences are technically viral, but widely carried by laboratory components, prone to result in false positives (14, 18). The viral functional cassettes of vectors are adopted from viruses. The inclusion of them in the reference database can raise a warning that if a query shows extremely high similarity with them, it should be concerned whether the sample is contaminated by exogenous false positives (18). Besides, attenuated viral strains are widely used in human and animal vaccinations to combat infectious diseases. It is important to distinguish them from field strains in clinic diagnosis. Vaccine sequences added here cover 15 attenuated viruses commonly used in humans and animals against mumps, Japanese encephalitis, equine infectious anemia and porcine epidemic diarrhea, etc.. By such augmentation, the database was finalized as eukaryotic viral reference database (EVRD), the nt and aa sequences were respectively archived in EVRD-nt and EVRD-aa branches. EVRD-nt has 558,673 sequences with average length of 2,943 bp covering 117 families, while EVRD-aa catalogs 1,256,089 sequences from 115 families with average length of 371 aa. EVRD-nt additionally records viroid sequences within the families *Avsunviroidae* and *Pospiviroidae*.

### EVRD improves the accuracy and efficiency of viromic analysis

The performance of EVRD was evaluated in viromic analysis by comparison of its ability to avoid false positives (accuracy), possibility to miss true viral contigs (coverage), and time to complete the analysis (efficiency) with Genbank (for nt) and UniProt (for aa) viral branches, and RVDB (v21.0) using nine viral metagenomic data of pigs, bats and humans. The results at the read level revealed that 13,417,025 reads in the nine datasets were annotated to be viruses by at least one of the databases, covering 47 families with 15 exclusively invisible to EVRD-nt in some datasets (Fig. 5). Majority (88.1%) of these virus-like reads (VLRs) were co-annotated by them, suggesting a high consistency using the three databases (Figs. 5 and 6A). Among those inconsistently annotated VLRs, 60.9% were exclusively annotated by RVDB-nt (subset R in Fig. 6A), followed by 38.2% being co-annotated by RVDB-nt and GenBank (G∩R in Fig. 6A).

**Fig 5.**
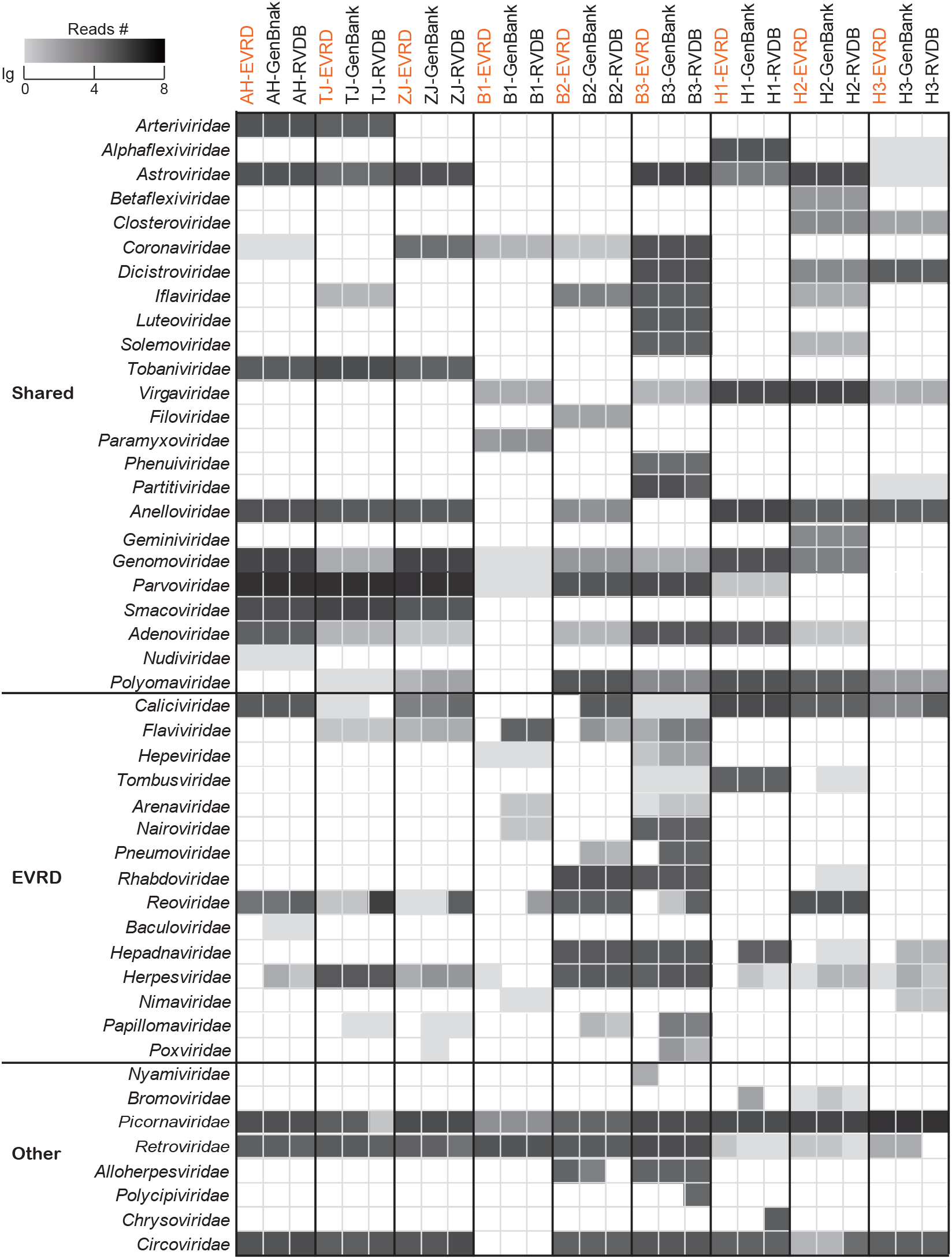
Comparison of the VLR numbers in nine viromic data sets annotated using blastn search against EVRD-nt (highlighted in orange), GenBank and RVDB-nt. Viral families are divided into parts of ‘Shared’, ‘EVRD’ and ‘Other’, corresponding to families that are co-annotated by the three reference databases, not annotated by EVRD in certain data sets, and annotated by one or two reference databases in certain data sets, respectively.

**Fig 6.**
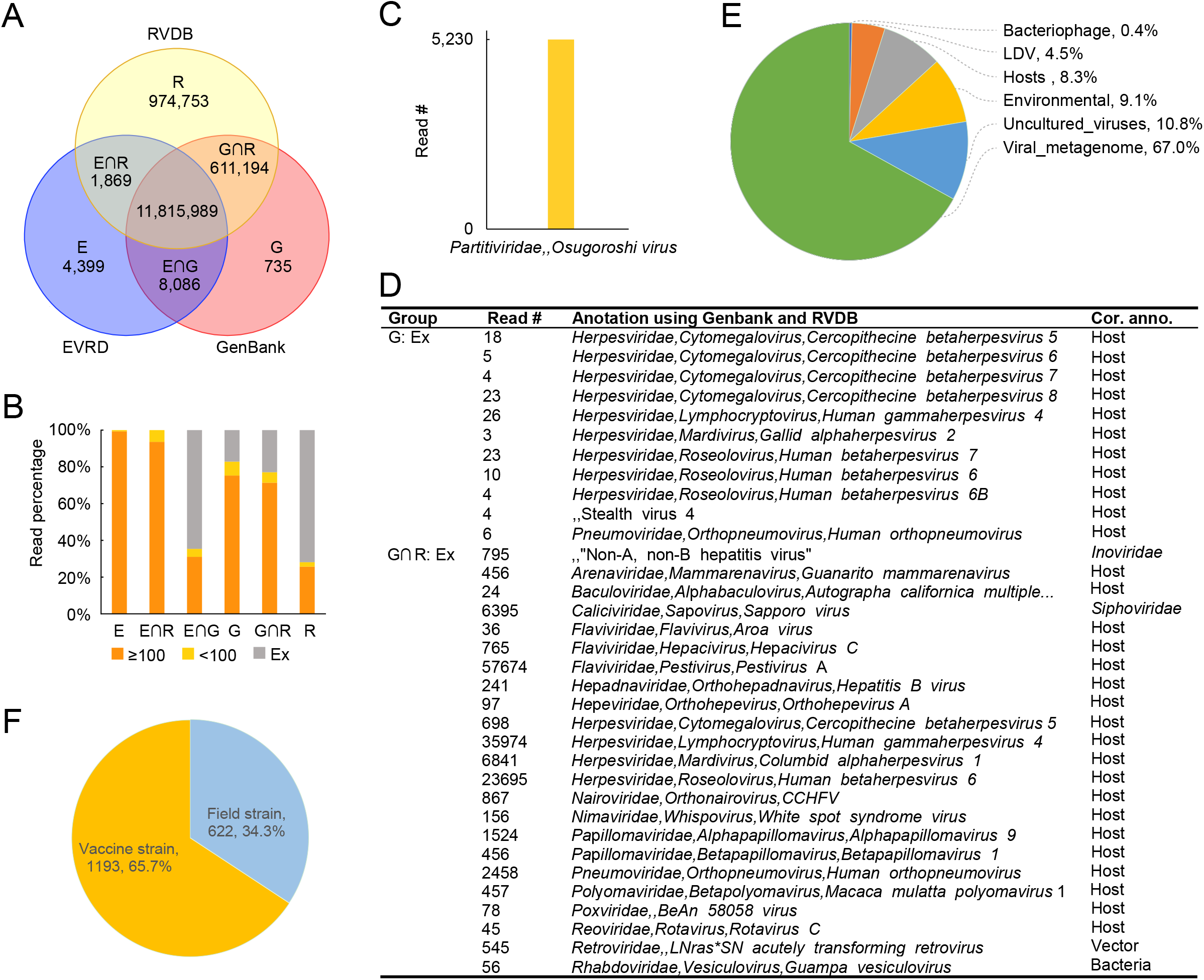
Identification of VLRs. A) VLRs were annotated using different databases with read numbers shown in each subset; B) The annotations of VLRs in the six subsets were improved using length cutoff 100 (orange bars), some VLRs can be annotated using the other databases with length < 100 (yellow bars), but there were still some VLRs (gray bars, labeled using Ex) unable to be annotated by the other databases even length was loosened to < 100; C) The Ex VLRs in subset E∩G were all related to Osugoroshi viruses within the family *Partitiviridae*; D) The Ex VLRs in subsets G and G∩R were all associated to HGSs; E) The Ex VLRs in subset R were predominantly annotated by RVDB-exclusive viral metagenomes; F) The PRRSV VLRs in data set AH belonged to vaccine and field strains based on the annotation using EVRD-nt.

The criterion used to determine whether a sequence is viral has a substantial impact on the annotation of these inconsistent reads. Some of these HTS datasets were generated with an insert size of 125 bp, so the requirement of alignment length ≥ 120 is a little stringent to them and has excluded many true positives. If we loosened the length cutoff to 100, such consistency was variably improved (Fig. 6B). Almost all of VLRs in subsets E and E∩R were annotated by the other database(s) using a loose length cutoff (Fig. 6B). But there were still lots of reads unable to be annotated by certain database(s) even using a loose length cutoff (illustrated using Ex in Fig. 6B). After improvement, 5,230 VLRs in E∩G remained unable to be annotated by RVDB-nt. All of these reads were related to Osugoroshi viruses within the family *Partitiviridae* that were recently released to the public by GenBank and have yet been synchronized in RVRD-nt v21.0 (Fig. 6C). The Ex VLRs in subsets G and G∩R, and their *de novo* assemblies, were all annotated to HGSs (Fig. 6D), i.e., they were false positives. The overwhelming majority (95.5%) of Ex VLRs in subsets R were related to sequences that are unrelated to eukaryotic viral pathogens and exclusively recruited by RVDB-nt, i.e., viral metagenomes, uncultured viruses, environmental samples, host-derived endogenous viral elements and bacteriophage (Fig. 6E). The remaining 4.5% were related to microorganism-infecting LDVs, such as pandora viruses and pithoviruses (Fig. 6E).

*de novo* assemblies (≥ 1000 bp) were also annotated using these databases. Compared to the results revealed using reads, 22 viral families were lost including *Filoviridae* that has proved to be present in samples (38). The annotation using EVRD-nt excluded the false positives of *Caliciviridae, Reoviridae* and *Herpesviridae* in certain datasets, indicating an improvement of accuracy at the contig level. Though the annotation using aa references of the three databases all showed higher specificity at the read and contig levels, EVRD-aa improved more significantly with exclusion of the false positives from *Reoviridae, Parvoviridae* and *Mitoviridae*, etc. These results indicated that the de-heterogeneity of our EVRD does not sacrifice the detection spectrum of eukaryotic viruses, rather significantly improves the specificity and accuracy of viromic annotation via reduction of erroneous annotation.

We did not find any viromic annotations tagged with ‘LCD’ or ‘Vector’, indicating no contamination of laboratory component- and vector-derived sequences in these datasets. But of special note is that, besides 622 reads in dataset AH annotated to porcine reproductive and respiratory syndrome virus (PRRSV) field strains, there were another 1,193 reads annotated to PRRSV vaccine strain in the dataset (Fig. 6F), indicating co-circulation of field viruses and vaccine strains in the farm, which should be especially concerning, since new viruses could be generated through recombination between field viruses and vaccine strains, resulting in vaccine failure (16). Viromic annotation is quite time- and computing resource-consuming. A small-scale reference database can shorten the analytic time and minimize the computing resource. With an entry-level platform, analyses of reads or contigs at nt or aa levels using EVRD were 1.8-3.3 and 1.9-3.2 times faster than using GenBank/UniProt and RVDB, respectively, indicating that EVRD is more efficient.

EVRD can be typically applied to, but not limited to, the virologic scenarios below. Accurate determination of causative agents is a priority in clinical diagnosis of viral diseases. However, the heterogeneity of reference database often produces confusing even wrong conclusion. Our previous viromic analyses often found sequences of CCHFV, HEV and BVDV, etc., but which were finally verified to be false positives. This phenomenon also occurred widely in other viromic studies (17, 18, 39, 40). For example, African swine fever virus was surprisingly found in a bat virome (40), which was highly unconvincing and most likely due to misannotation of host sequence, since African swine fever virus is particularly host-specific and only infects swine (41). EVRD has deleted the disturbing HGSs in reference sequences, thus reduces such confusion by preventing misannotation at source. EVRD can also improve the taxonomic classification of viral sequences in assessment of virus diversity (26). In such analysis, viral contigs need to be clustered with reference sequences, but the HGSs, especially the cross-family misclassified ones, will disturb the boundary of virus clusters, even result in incorrect taxonomic classification. In addition, multiple sequence alignment (MSA) is prone to be corrupted by HGSs, the refined EVRD sequences can help build high-quality MSAs that are basis of profiles of clustered sequences (not included in this study), thus favoring the exploration of remote viruses.

Critical is to correctly annotate sequences in viral disease diagnosis and viromic analysis. Besides utilizing a high-quality reference database, other measures can be taken into account. First, reasonable bioinformatic pipelines should be implemented for different purposes. Annotation using reads provides richer information than using contigs, especially for ultra-low abundant viruses (38, 42), hence could be considered in viral disease diagnosis. But sequence completeness is a priority in viral ecology, thus assembly is preferentially performed before annotation (26). Second, criterion to determine a viral sequence has non-ignorable impact on annotation. As to reads, criterion is mainly based on evalue, but the alignment length is also an important factor to help increase the confidence level of annotation. Besides evalue and length, the requirement of a minimum of gene number has been widely considered in contig annotation (26). Third, the quality of assemblies should be seriously considered in contig annotation. There are many means to improve assembly quality, such as choosing a suitable software (43), employing a rational sample treatment protocol (44), reducing the bias induced by random amplification (45). A classification of host and other microorganism reads prior to *de novo* assembly could help reduce chimeric contigs. Fourth, of special note is the annotation of remote viruses. Due to lack of enough known references, it is often difficult to precisely annotate these contigs based on similarity search. A combination of multiple advanced annotations, such as profile-based classification and deep learning-based recognition, is permissive and necessary (46-48). Last but not least, a final check provides an additional guarantee for high-quality annotation (49). Host contamination should be eliminated as much as possible. Prokaryotic contamination can be determined using CheckV, but a different strategy is needed to deal with eukaryotic contamination (49, 50). Contigs with extraordinary genomic structure and/or organization, e.g., excessive length and long noncoding region, might be resultants of misassembly or insertion of exotic sequences, and should be further verified. In conclusion, in order to control contamination at source, sequences with their annotations should be carefully inspected by submitter before submitting to public databases.

When using EVRD, users need to take note of several aspects. We excluded LDVs infecting eukaryotic microorganisms, due to their extraordinarily large and complicated genomes and lacking evidence to cause diseases in vertebrates (51-53). Though we have deleted hundreds of HGSs of vertebrate LDVs from families like *Herpesviridae, Poxviridae*, there are still some ambiguous sequences that can be treated as host HGSs if using loose criteria. Those viruses, along with retroviruses, can exchange genomic fragments with hosts, and have undergone long-term co-evolution with host, which would smooth the distinctive trait of those sequences between viruses and hosts (19, 20, 54). Thus, annotations to these viruses using EVRD should still be verified with caution. Additionally, these tagged warning sequences in EVRD are very useful, but they are just partial and only represent the sequences we have searched so far. We will keep the database updated with new advances in this regard.

## Conclusion

A high-quality virus reference database is critical to accurate analysis of viral sequences. In this study an improved reference database of eukaryotic viruses has been built from existing public GenBank/UniProt databases based on a stringent scrutiny pipeline to remove hundreds of confusing HGSs. It showed better accuracy and efficiency in annotation of eukaryotic viromes compared to its parent databases and the extensive RVDB. With functional augmentation using tagged risk and vaccine viruses, EVRD significantly facilitates the genomic analyses in applications like viral disease diagnosis, taxonomic classification, and new virus detection and identification.

## Methods

### Heterogeneity scrutiny pipeline for nucleotide sequences

#### I) Preliminary filtration

We first generated the taxonomic lineages of all sequences, then removed those lineages infecting bacteria, archaea, fungi and eukaryotic microorganisms using the relationship of virus and host recorded in ViralZone database (55). In addition, there are a large number of sequences that cannot be assigned to a complete lineage, we searched their definition using keywords and removed the sequences related to prokaryotic and environmental viruses and metagenomes, such as bacteriophage/phage, environment, uncultured and ameba. Division gbvrl also deposits numerous sequences ≤200 bp, which are highly similar to these longer sequences, and contribute a little to diagnosis and virus identification, hence were also removed.

#### II) Host genome scrutiny

In this part, fragments of host genomes in the remaining sequences of PDS were scrutinized. Genomic assemblies of human (n=1), pig (n=1), bats (n=7), rodents (n=2), arthropods (n=11), cattle (n=1), dog (n=1), cat (n=1), sheep (n=1), chicken (n=1) and mallard (n=1) were used to BLASTn search against these sequences with a maximum of 1000 subjects to show alignments (length ≥ 150 and identity ≥ 85%). Retroviruses can infect almost all vertebrates, resulting in thousands of loci of retroviral sequences in vertebrate genomes (54). Here we did not challenge the known ambiguity of retroviruses, hence hits to retroviruses were not considered. The aligned sequences of subject were extracted and subjected to blastn search against nt database to further validate their identities. The top 100 hits of each sequences were kept and, within which, if ≥80% hits were annotated to nonviral, the aligned sequence was considered heterogenous. The original sequence was removed from PDS if its heterogenous part comprises ≥80% of its length, or trimmed by deleting the heterogenous parts, such threshold was also applied to the following treatments. The rest of PDS was subjected to a next round of scrutiny until no host genomic fragments were found.

#### III) Vector sequence scrutiny

To detect HGSs derived from backbones or functional cassettes of vectors, UniVec database and sequences ≥1,000 bp under the GenBank taxonomy of vectors (uid: 29278) were downloaded. As vectors have many functional cassettes originated from viruses, such as SV40 and CMV promoters, retroviral *gag* and *pol* elements, these vector-originating HGSs in PDS were carefully detected and examined using the following procedure to prevent any erroneous deletions of genuine viral sequences. First, we generated a non-viral protein core (NVPC) that consists of nonviral expression elements (n=13,287) born in vectors. To achieve that, those protein sequences ≥ 100 aa encoded by vectors were de-replicated using cd-hit v4.8.1 with 99% similarity at 90% coverage for the shorter sequences (56). The resulting representatives (n=17,236) were blastp searched against the nr database using Diamond with maximum number of 100 target sequences to report alignments (57). The representatives classified as viruses using a majority-rules approach were discarded, while the rest (n=15,220) were further queried against the UniProt viruses branch. These unaligned sequences (n=12,603) were technically nonviral and classified into NVPC, while these aligned (n=2,617) were manually inspected by online blastx search against nr database with these (n=684) annotated to nonviral products being classified into NVPC. Sequences in PDS were blastx searched against NVPC using Diamond with these showing ≥99% similarity over alignment ≥60 aa with subjects being pruned. In addition, UniVec was used to identify adapters, linkers, and primers often used to clone sequences. The remaining sequences in PDS were further scrutinized using procedure introduced in part II with the same criteria. Briefly, these vector sequences were used as query to search possible subjects in PDS using blastn. Hits in PDS were further validated by blastn search against nt database. After removal of those vector-originated sequences, the rest of PDS were examined by another round of scrutiny until no vector sequences exist.

#### IV) Annotation cross scrutiny

Erroneously taxonomic annotation of viral sequences was detected by all-against-all blastn search with a maximum of 1000 subjects to show alignments. We found that there are a large number of sequences with correct taxonomic annotation showing intra-family cross-species/genus blastn hits, such as *Betacoronavirus*/*Gammacoronavirus* within the family *Coronaviridae, Tetraparvovirus*/*Protoparvovirus* of the family *Parvovridae*, and *Circovirus*/*Cyclovirus* within the family *Circoviridae*, which were likely ascribed to high similarity between species/genus. Hence, we inspected annotation at the family level. Here we defined that a blastn hit is significant if its e-value is ≤ 1e-50 and length ≥500. If the proportion of alignments that were generated by a query against subjects of different family to all alignments of the query is ≥80%, the query was considered being possibly misclassified, which was further subjected to genomic organization identification, in which if the genomic organization of the query is not of typical feature its defined taxonomic lineage should have, the query was truly misclassified and removed from PDS. During treatment, we noted that some sequences had a few alignments (usually ≤10) that show ≤80% similarities with subjects of different family, we kept their original annotations since lack of enough references in GenBank to determine their true taxonomic lineages.

#### V) Cross check of viral metagenomes

Previous study showed that some contaminant viral sequences are highly prevalent in cross-host HTS-based viromic data, which might be linked to biological or synthetic products (18). To examine whether cross-host sequences exist in database, the remaining sequences in PDS were subjected to cross check of viral metagenomes. A total of 15 viromic raw data sets covering human, bat, tick, rodent, bovine, pig and avian were downloaded from SRA and respectively *de novo* assembled. Contigs ≥ 1000 bp were subjected to blastn search against PDS with a maximum of 1000 subjects to show alignments. If a subject was matched by contigs from viromic data sets of ≥ two different hosts with alignment ≥ 150 bp and identity ≥ 80%, it was classified as suspicious sequence and further validated by blastn search against nt database. If a suspicious sequence was annotated to non-viral species by blastn search against nt database, it was considered as a truly exogenous contaminating sequence and removed from PDS. However, if a suspicious sequence was still annotated to virus and shared 99% nt identities with viromic contigs of ≥ two different hosts, it was considered as a truly viral sequence but probably originated from laboratory-component derived viral sequence contamination, hence was retained in PDS but was tagged as LCD. The remaining suspicious sequences were passed and kept in PDS.

### Heterogeneity scrutiny pipeline for viral protein sequences

The protein sequences retrieved from UniProt virus division were subjected to scrutiny as described above with minor modification. We first checked their representativeness. In case there are any coding regions not annotated by the original submitters, all proteins of PDS nt sequences prior to filtration were *de novo* predicted using prodigal v2.6.3 with meta mode. Proteins ≥ 50 aa were blastp searched against UniProt viral division (evalue ≤ 1e-10 and pident ≥ 90), and results revealed that UniProt viral division has high representativeness with 99.6% consistency to the prediction of GenBank viruses. In the step of preliminary filtration, we removed those non-eukaryotic viral sequences and those ≤ 30 aa. The remaining sequences were used to blastp search against the genomic protein sequences of the hosts to detect any potential host contaminants (length ≥ 100 and identity ≥ 90%), these host contaminants if detected were further subjected to blastp search against nr database to finally identify whether they are host protein sequences with the same criterion used in nt identification. The scrutiny was iteratively performed until no host contaminants were found. In the vector sequence scrutiny, a blastp search of PDS against NVPC was conducted to find any vector contaminants. The queries with identity ≥90% over alignment ≥100 with NVPC were further validated and treated as described in host protein scrutiny. The annotation cross scrutiny of viral protein sequences was nearly the same as that in nt scrutiny but only that the all-against-all blastp hits were considered significant if their e-values were ≤1e-50 and length ≥100. In cross check of the viral metagenomes, contigs ≥1000 bp were subjected to blastx search against viral protein sequences. The viral protein sequences were considered suspicious if they matched to contigs of viral metagenomes from ≥ two host species, and subjected to further validated by blastp search against nr database as described in cross check of the viral metagenomes.

### EVRD finalization

After above scrutiny, the sequences in PDS are still very redundant, hence a de-redundance procedure is applied to downsize PDS. Clustering of viral nt and aa sequences was performed using MMseq2 (58) with sequence similarity threshold of 0.99 and 90% coverage of the short sequence. Viral sequences if identified as LCD with real virus origin (14, 18) are tagged by ‘LCD’ as risk sequences before adding into PDS. To better distinguish viral functional cassettes from true virus sequences, the sequences corresponding to the regulatory classes of promoter, terminator and enhancer, and/or the notes containing the word of ‘virus’ were extracted from vectors, and subjected to blastn search against the non-redundant PDS, the sequences verified to be viral were de-replicated and also added to PDS with the tag ‘Vector’. In addition, we collected vaccine strains commonly used in humans and animals such as pigs, chickens and dogs, via searching in publications or by personal communication. These vaccine nt sequences were also added in PDS with the tag ‘Vaccine’.

### Performance evaluation of EVRD

Nine viral metagenomic data sets were first subjected to host genome removal using Bowtie2 (v2.4.1) with sensitive mode, and then taxonomically classified using Kraken2 (v2.0.9-beta) to remove bacterial, archaeal and fungal reads. The unassigned reads were firstly blastn (evalue ≤ 10e-5 and length ≥ 120) and blastx (evalue ≤ 10e-5 and length ≥ 40) searched against these databases. Then they were *de novo* assembled using megahit (v1.2.9). Contigs ≥ 1000 bp were retained for blastn (v2.10.0) and diamond blastx (v0.9.35) search against nt and aa reference databases, respectively. The blastn hit of a contig to a subject with one alignment of evalue ≤ 10e-10 and length ≥ 450 or ≥ two alignments of evalue ≤ 10e-5 and length ≥ 150 was considered positive, and the blastx hit to a subject was recognized positive if it had one alignment of evalue ≤ 10e-10 and length ≥ 150 or ≥ two alignments of evalue ≤ 10e-5 and length ≥50. The positive reads and contigs were further verified by blastn/x search against nt/nr databases (16). All blast searches were performed using 12 x86_64 CPUs of an Inter® Xeon® Gold 2.660 GHz processor. To detect waring sequences tagged by “LCD”, “Vector” and “Vaccine” in the viromic annotation using EVRD, we defined a rigorous cutoff, i.e., a sequence with positive blastn hit to a tagged subject with identity ≥99% and coverage of the query ≥90% was considered risk and vaccine sequence.

## Availability of data and materials

All data used here were downloaded from relevant databases. The key intermediate data (NVPC) and essential codes are available from http://github.com/BH-Lab/EVRD. EVRD reported here (the first release: 2021.03) is based on the viral branches (version 2021.03) of Genbank and UniProt, and is scheduled to annual update, which is freely accessible at http://cvri.caas.cn/kxyj/yjfx/bfdb/EVRD.index.htm.

## Competing interests

The authors declare that they have no conflict of interest

## Notes

https://cvri.caas.cn/kxyj/yjfx/bfbd/EVRD/index.htm

### Competing Interest Statement

The authors have declared no competing interest.

